# Nicotinamide Metabolism Reprogramming Drives Reversible Senescence of Glioblastoma Cells

**DOI:** 10.1101/2024.10.18.619131

**Authors:** Ashwin Narayanan, Mirca S Saurty-Seerunghen, Jessica Michieletto, Virgile Delaunay, Arnaud Bruneel, Thierry Dupré, Chris Ottolenghi, Clément Pontoizeau, Lucrezia Ciccone, Andreas De La Vara, Ahmed Idbaih, Laurent Turchi, Thierry Virolle, Hervé Chneiweiss, Marie-Pierre Junier, Elias A. El-Habr

**Author notes:** Corresponding author: 7 quai Saint-Bernard 75005, Paris, France. co-first author (Ashwin Narayanan, Mirca S Saurty-Seerunghen). co-last authors (Marie-Pierre Junier, Elias A. El-Habr).

## Abstract

Recent studies show that metabolites, beyond their metabolic roles, can induce significant changes in cell behavior. Herein, we investigate the non-canonical role of nicotinamide (vitamin B3) on glioblastoma (GB) cell behavior. Nicotinamide induced senescence in GB cells, characterized by reduced proliferation, chromatin reorganization, increased DNA damage, enhanced beta-galactosidase activity, and decreased Lamin B1 expression. Nicotinamide-induced senescence was accompanied by an unexpected reprogramming of its metabolism, marked by simultaneous downregulated transcription of NNMT (nicotinamide N-methyltransferase) and NAMPT (nicotinamide phosphoribosyl-transferase). Nicotinamide effects on GB cells were mediated by decreased levels of SOX2. Consistently, analyses of patients’ single cell transcriptome datasets showed that GB cells with low NNMT and NAMPT expression levels were enriched in gene modules related to senescence. Remarkably, senescent GB cells retained tumor-forming ability *in vivo*, albeit to a lesser extent compared to control cells. Further experiments at the single-cell level and transcriptomic analyses demonstrated that nicotinamide-induced senescence in GB cells is fully reversible. Overall, our findings identify a novel reversible senescent state in GB tumors and highlight the non-canonical role of NAM as a key driver of cancer cell plasticity.

## Introduction

Changes in the functional state of both normal and cancerous cells are systematically accompanied by metabolic reprogramming. Initially viewed as a passive adaptation to meet energy demands and support biomass production, recent evidence challenges this view, revealing that metabolic reprogramming actively drives changes in cell behavior. Key cellular processes, such as epigenetic regulations, are directly influenced by specific metabolites (Reid et al., 2017; Tsogtbaatar et al., 2020). Our results, along with others, show that fluctuations in intracellular levels of several metabolites -serving as substrates, cofactors, or inhibitors of epigenetic enzymes- directly influence gene expression, thereby inducing changes in cell functional states (Carey et al., 2015; El-Habr et al., 2017; Liu and Wellen, 2020).

One of these metabolites, the universal methyl donor S-Adenosyl methionine (SAM), exerts significant influence on cell functional states by modulating histone and DNA methylation. Nicotinamide N-methyltransferase (NNMT) is a key regulator of SAM levels. Using SAM as a methyl donor, NNMT catalyzes the methylation of nicotinamide (NAM), the inactive form of vitamin B3, yielding to the formation of S-adenosyl-L-homocysteine (SAH) and 1-methylnicotinamide (MNAM) (Pissios, 2017). Overexpression of NNMT depletes intracellular SAM, reducing its availability for other methyltransferases involved in DNA and histone methylation. This mechanism is instrumental in the maintenance of naïve embryonic stem cell pluripotency, which relies on low levels of DNA and histone methylation (Sperber et al., 2015). NNMT overexpression is also observed in various cancer types, including glioblastoma (GB) (Thirant et al., 2012), the most aggressive primary brain tumor, where it contributes to epigenetic alterations that drive tumor aggressiveness (Jung et al., 2017; Palanichamy et al., 2017).

Beyond SAM, high NNMT expression levels are expected to decrease NAM availability. The other enzyme consuming NAM is nicotinamide phosphoribosyltransferase (NAMPT), which catalyzes the first rate-limiting step of NAD^+^ biosynthesis. NAD^+^ plays crucial roles in redox signaling implicated in cellular energy production and acts as cofactor for non-redox enzymes like sirtuins, which mediate deacetylation of histone and non-histone proteins. Like NNMT, NAMPT is overexpressed in GB, leading to increased NAD^+^ production, which enhances GB cell aggressiveness (Gujar et al., 2016; Jung et al., 2017; Lucena-Cacace et al., 2019). Despite accumulating evidence highlighting the contribution of NNMT and NAMPT to GB growth, the direct role of NAM itself in modulating GB cell behavior remains unexplored.

Here, we show that NAM accumulation in several GB patient-derived cell lines (GB-PDC) induced their transition to senescence. NAM-induced senescence of GB-PDC was accompanied by down-regulated NNMT and NAMPT expression. Analysis of single-cell transcriptomic datasets derived from GB patients confirmed that malignant cells with low NNMT and NAMPT expression are enriched in gene modules associated with senescence. Further investigations, including *in vivo* experiments, single-cell and transcriptomic analyses, revealed the reversible nature of NAM-induced senescence in GB cells. Collectively, our results highlight a novel, reversible senescent state in GB cells, controlled by a single metabolite, NAM, offering new insights into GB cell plasticity.

## Results

### NNMT downregulation during loss of GB-PDC tumorigenic properties is accompanied by accumulation of NAM

We utilized an aggressive GB-PDC model (TG1) and its counterpart expressing miR- 302-367 (TG1-miR), which has lost its tumorigenic properties (Fareh et al., 2012), to compare NNMT expression levels. Our analysis demonstrated a marked reduction in both NNMT mRNA and protein levels in TG1-miR cells compared to the TG1 (Fig. S1A and B). To assess the effect of NNMT downregulation on the intracellular levels of its substrates and products, we performed gas chromatography/mass spectrometry (GC/MS) and liquid chromatography/ MS/MS, as previously detailed (El-Habr et al., 2017). Our analysis revealed a significant decrease in NNMT products, MNAM and SAH (Fig. 1A). Interestingly, while NNMT downregulation did not affect SAM levels, it led to an accumulation of intracellular NAM (Fig. 1A). These findings suggest that elevated intracellular NAM levels might mediate the effects observed during NNMT downregulation and the subsequent loss of GB-PDC tumorigenicity. To test this hypothesis, we examined the effects of NAM on GB-PDC properties.

**Figure 1.**
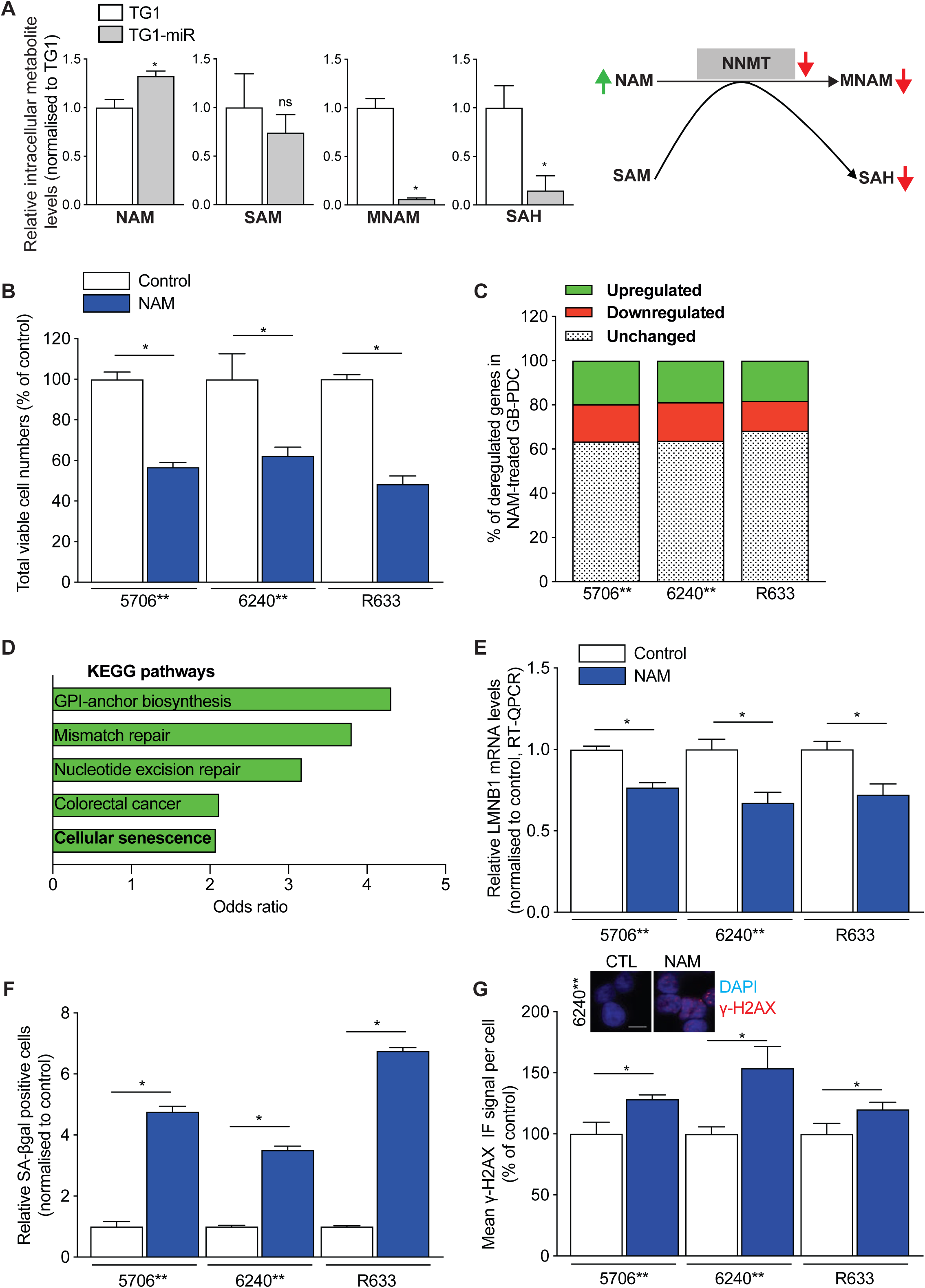
NAM induces senescence of GB-PDC. **A**. Increased NAM levels accompany NNMT downregulation observed in TG1-miR. Schematic representation of the enzymatic reaction catalyzed by NNMT, with graphs showing the levels of each of its components. Green and red color indicates increased and decreased metabolites/enzyme respectively in TG1-miR compared to TG1. Metabolite intracellular levels normalized to TG1. Unpaired *t* test with Welch’s correction, mean ± SD, n = 3 independent biological samples (SAM and SAH) or n = 6 (NAM and MNAM). SAM, S-Adenosyl methionine; SAH, S-adenosyl-L-homocysteine; MNAM, 1-methylnicotinamide. **B**. NAM inhibits cell growth of GB-PDC. Total numbers of viable cells evaluated after 1- or 2-weeks treatment. Unpaired *t* test with Welch’s correction, mean ± SD, n = 3 independent biological samples. **C**. NAM alters GB-PDC transcriptomic profiles. Gene fractions with upregulated, downregulated or unchanged expression following NAM treatment are highlighted in green, red, or with a dot motif, respectively. **D**. Pathway enrichment analysis (PEA) on gene differentially expressed highlights senescence in NAM-treated GB-PDC. Top 5 KEGG terms of PEA done on the upregulated genes shared across all GB-PDC are shown. Remaining terms and further analyses are listed in Table S1. **E**. NAM decreases LMNB1 mRNA levels of GB-PDC. RT-QPCR assays, unpaired *t* test with Welch’s correction, mean ± SD, n = 3 independent biological samples. **F**. NAM increases beta-galactosidase activity of GB-PDC. Quantification of beta-galactosidase (SA-βgal) positive cells, unpaired *t* test with Welch’s correction, mean ± SD, n = 3 independent biological samples. **G**. NAM increases *γ-* H2AX levels of GB-PDC. Quantification of *γ-*H2AX (phosphorylated H2AX) immunofluorescent (IF) signal. Unpaired *t* test with Welch’s correction, mean ± SD, n = 9 independent biological samples. Images illustrate an example of *γ-*H2AX immunocytochemical detection in control (CTL) and NAM-treated (NAM) 6240** GB-PDC. Scale bar 10 µm. Statistical significance: ns non-significant, * P ≤ 0.05.

### NAM induce senescence features in GB-PDC

We assessed the effects of NAM on three distinct adult GB-PDC (5706**, 6240**, and R633) (El-Habr et al., 2017), maintained in EGF/bFGF supplemented media that preserves their tumorigenic properties. Measure of NAD(P)^+^/NAD(P)H ratios with a WST-1 assay in GB-PDC treated with 10 to 50 mM NAM showed a significant decrease in their metabolic activity in response to 30 mM NAM (Fig. S1C). Intracellular accumulation of NAM was confirmed by HPLC (Fig. S1D). Further experiments were conducted using this NAM concentration. FACS analysis showed that NAM treatment did not affect cell viability (Fig. S1E). However, it significantly inhibited their proliferation, reducing cell numbers by 40-50%, as shown by cell counting and Trypan blue exclusion assays (Fig. 1B). To understand the molecular mechanisms underlying NAM-induced inhibition of GB-PDC proliferation, we conducted RNA sequencing to compare the transcriptomic profiles of NAM-treated and control cells. NAM treatment induced deregulated expression of 32 to 37% of detected genes (Fig. 1C, Tables S1). These results indicate that NAM induces a cytostatic effect on GB-PDC, accompanied by a broad transcriptomic deregulation.

To elucidate the molecular pathways underlying NAM-induced inhibition of GB-PDC proliferation, we conducted pathway enrichment analysis on the deregulated genes shared across all GB-PDC (Tables S1). Pathway enrichment analysis of both deregulated and specifically upregulated genes identified “Cellular Senescence” as a significantly enriched biological process across all GB-PDCs (Fig. 1D, Table S1). Senescence represents a multifaceted cell state characterized by cell cycle arrest, morphological alterations, chromatin reorganization, and modified gene expression. This state can be attained via various molecular pathways, with senescent cells exhibiting several molecular and phenotypic changes that serve as markers (Hernandez-Segura et al., 2018). Since senescence lacks a universal biomarker (Hernandez-Segura et al., 2018), we assessed whether NAM treatment induced a senescent state in GB-PDCs by evaluating multiple well-established senescence markers.

Nuclear lamina, crucial for maintaining nuclear morphological integrity, primarily consists of Lamin B1, which is commonly downregulated during senescence (Freund et al., 2012). In our study, NAM reduced transcript levels of LMNB1, encoding Lamin B1, as observed by RT-QPCR and RNA-sequencing (Fig. 1E, Table S1). Another hallmark of senescent cells is increased lysosomal content, leading to enhanced activity of lysosomal enzyme senescence-associated beta-galactosidase (SA-βgal) (Lee et al., 2006). We observed a robust increase (3.5- to 7-fold) in the number of SA-βgal-positive cells upon NAM treatment of GB-PDC compared to controls (Fig. 1F). Senescent cells also display morphological alterations (Sikora et al., 2016), and FACS analysis confirmed altered cell morphology after NAM treatment (Fig. S1F). Finally, the pathway analysis also revealed an enrichment of “Mismatch repair” genes (Fig. 1D, Table S1). DNA damage being another hallmark of senescence (Herranz and Gil, 2018), we thus assessed the impact of NAM on phosphorylation levels of H2AX (γ-H2AX), a DNA damage marker (Fumagalli et al., 2012).

Immunofluorescent imaging and Western blot analysis revealed higher γ-H2AX levels in NAM-treated GB-PDC compared to controls (Fig. 1G, S1G).

These findings collectively demonstrate that NAM induces a shift of GB-PDC towards a cell state with senescent features.

### NAM inhibits NNMT transcript levels via SOX2 repression

NAM-induced senescence of GB-PDC was accompanied by a drastic reduction of the expression of the two NAM consuming enzymes, NNMT and NAMPT, as revealed by RNA-sequencing, and confirmed by RT-QPCR and Western blot analysis (Fig. 2A, S2A, Table S1). These findings underscore that NAM-induced senescence in GB-PDC is accompanied by an unexpected reprogramming of NAM metabolism.

**Figure 2.**
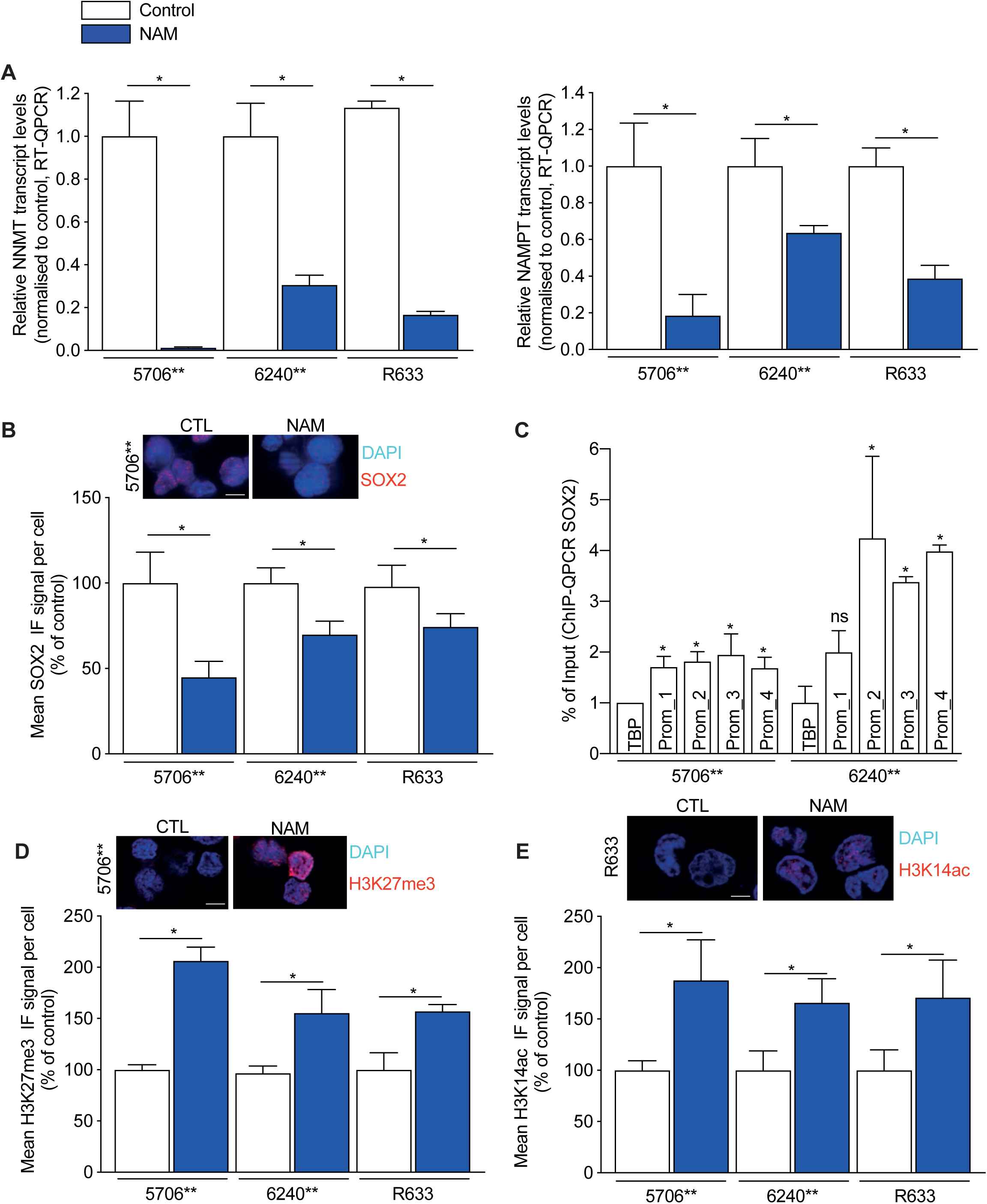
NAM induces reprogramming of its own metabolism by repressing SOX2, resulting in global epigenetic changes. **A.** NAM inhibits NNMT and NAMPT expression. Decreased NNMT (left panel) and NAMPT (right panel) mRNA levels in NAM-treated GB-PDC compared to control. RT- QPCR assays, unpaired *t* test with Welch’s correction, mean ± SD, n = 3 independent biological samples. **B**. Decreased SOX2 protein levels in NAM-treated GB-PDC. Quantification of SOX2 IF signal per cell. Unpaired *t* test with Welch’s correction, mean ± SD, n = 9 independent biological samples. Images illustrate an example of SOX2 immunocytochemical detection in control (CTL) and NAM-treated (NAM) 5706** GB-PDC. Scale bar 10 µm. **C**. ChIP-QPCR results showing enrichment in SOX2 binding sites in NNMT promoter, as compared to TBP. Mean ± SD, n = 3 independent biological samples. One way ANOVA. **D**. Increase in H3K27me3 levels in NAM-treated GB-PDC. Quantification of H3K27me3 IF signal per cell. Unpaired *t* test with Welch’s correction, mean ± SD, n = 9 independent biological samples. Images illustrate an example of H3K27me3 immunocytochemical detection in control (CTL) and NAM-treated (NAM) 5706** GB-PDC. Scale bar 10 µm. **E**. Increase in H3K14ac levels in NAM-treated GB-PDC. Quantification of H3K14ac IF signal per cell. Unpaired *t* test with Welch’s correction, mean ± SD, n = 9 independent biological samples. Images illustrate an example of H3K14ac immunocytochemical detection in control (CTL) and NAM-treated (NAM) 5706** GB-PDC. Scale bar 10 µm. Statistical significance: ns non-significant, * P ≤ 0.05.

We hypothesized that the transcription factor SOX2 mediates this reprogramming, given its role in controlling GB cell tumorigenicity (Bogeas et al., 2018; Garros-Regulez et al., 2016b; Suvà et al., 2014) and the observation that its downregulation leads to GB-PDC senescence (De Lope et al., 2019; Garros-Regulez et al., 2016a; Vinchure et al., 2021; Wu et al., 2020). Additionally, SOX2 acetylation has been shown to lead to its nuclear exclusion and subsequent degradation (Baltus et al., 2009; Yoon et al., 2014), and increased intracellular NAM levels have been reported to inhibit sirtuin deacetylases (Hwang and Song, 2017). We thus postulated that elevated NAM levels would reduce SOX2 protein levels. Immunofluorescent imaging confirmed a decrease of SOX2 nuclear signal in NAM-treated GB-PDC (Fig. 2B), in face of unchanged SOX2 transcript levels (Fig. S2B). SOX2 has been reported to bind to regulatory regions of NNMT gene (Lachmann et al., 2010), suggesting it may regulate NNMT expression. Consistent with this, we detected with ChIP-QPCR enrichment in several SOX2 binding sites within the NNMT promoter (Fig. 2C). These results indicate that SOX2 downregulation is a key mediator of the repressive effects of NAM on NNMT transcript levels.

### Global epigenetic modifications accompany NAM-induced senescence of GB-PDC

The significant transcriptome alterations in NAM-treated GB-PDC suggest global chromatin remodeling, a well-known hallmark of cellular senescence (Sun et al., 2018). NAM likely modulates DNA and histone methylation by repressing NNMT, which in turn increases SAM availability for other methyltransferases, potentially enhancing global methylation patterns. NAM-treated GB-PDC also showed increased expression of the methyltransferase EZH1 (Table S1, Fig. S2C), the catalytic subunit of Polycomb Repressive Complex 2 (PRC2), which catalyzes the trimethylation of histone H3 at lysine 27 (H3K27me3) (Piunti and Shilatifard, 2021). Consistently, immunofluorescent imaging demonstrated elevated global levels of H3K27me3 in NAM-treated GB-PDC compared to controls (Fig. 2D). In addition to histone methylation, NAM can also affect histone acetylation through inhibition of sirtuin deacetylases (Hwang and Song, 2017). Immunofluorescent imaging revealed a uniform increase in H3K14 acetylation, a marker associated with active transcription, across all NAM-treated GB-PDC (Fig. 2E), while H3K27 and H3K9 acetylation levels showed variable changes (Fig. S2D, S2E).

Collectively, these results demonstrate that global epigenetic reprogramming, a hallmark of senescence, is a key feature of NAM-induced GB-PDC senescence.

### GB patient cells with low NNMT and NAMPT expression show enrichment in genes associated with senescence

To confirm the relevance of our findings in the actual context of GB patients, we analyzed the distribution of NNMT and NAMPT expression across malignant cells in patient tumors. We utilized four publicly available single-cell RNA-sequencing (scRNA-seq) datasets from surgical resections of IDH-wildtype GB patients with diverse genomic abnormalities (Table S5) (Neftel et al., 2019; L. Wang et al., 2022; Yu et al., 2020). These datasets, derived from 65 tumors, comprise a total of 58,894 GB cells. Since null RNA counts obtained with scRNA-seq cannot be unambiguously attributed to an actual lack of gene expression rather than to a technical failure, we included in the analyses only cells with detectable expression of NNMT and NAMPT.

We found a heterogeneous distribution of NNMT and NAMPT transcript levels across malignant cells, indicating a variable activation of NAM metabolism within patient tumors (Fig. 3A, S3). We analyzed differentially expressed genes between cells with low and high NNMT and NAMPT expression by dividing them into two groups based on the mean expression of these two genes (Table S2). Pathway enrichment analysis of all deregulated, as well as upregulated genes in each dataset independently identified "Cellular Senescence" as significantly enriched in the NNMT^LOW^NAMPT^LOW^ cell group (Table S3, Fig. 3B). These results demonstrate that altered NAM metabolism distinguishes GB cells with senescent features within patient tumors, like our *in vitro* observations.

**Figure 3.**
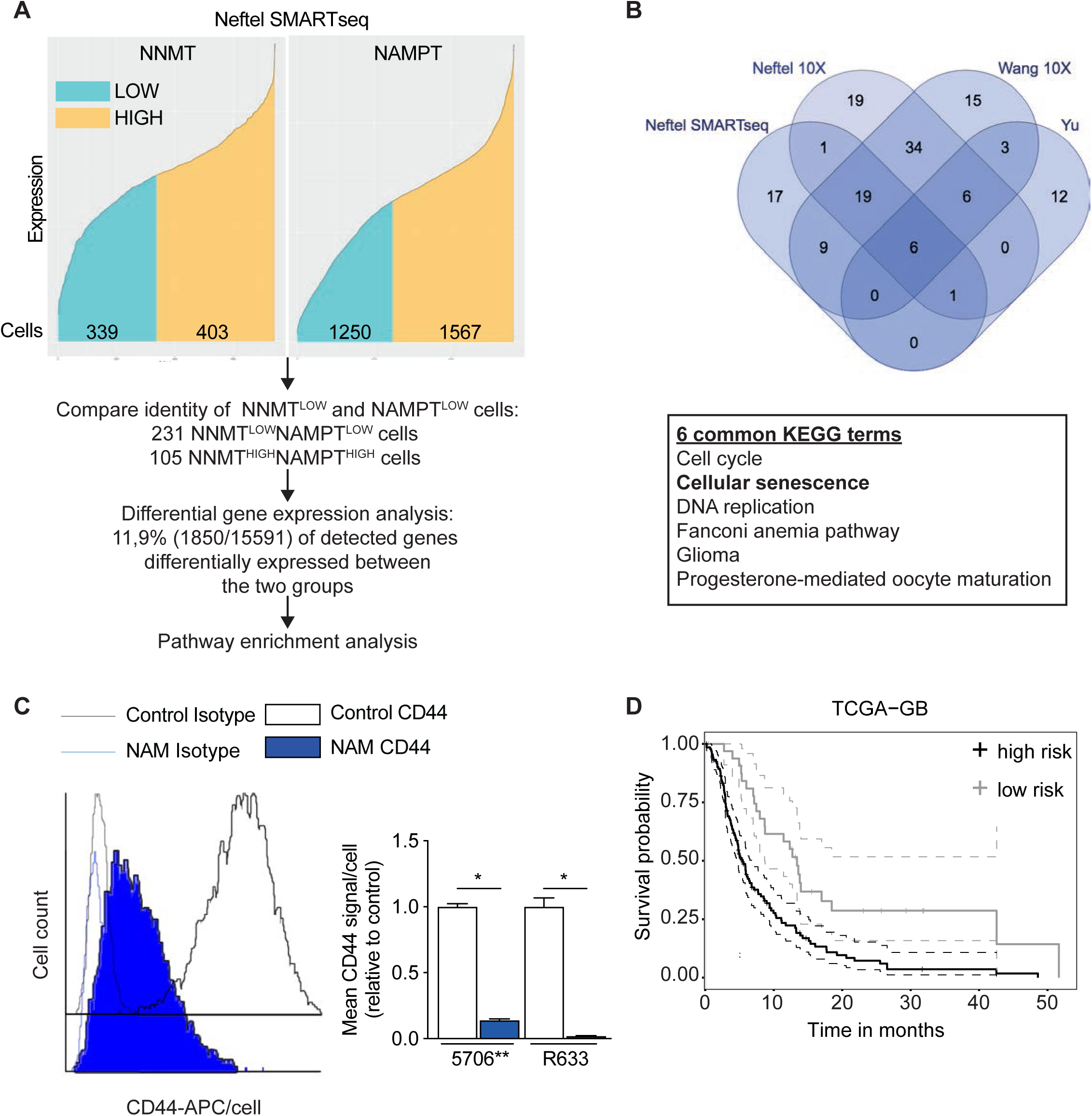
NAM metabolism reprogramming discriminates GB cells with senescent features within patient’s tissue. **A**. Scheme of the computational analysis of Neftel SMARTseq dataset. Cells were split into two groups, NNMT^LOW^NAMPT^LOW^ and NNMT^HIGH^NAMPT^HIGH^, based on the mean expression of NNMT and NAMPT genes. 231 and 105 cells were respectively retained as NNMT^LOW^NAMPT^LOW^ and NNMT^HIGH^NAMPT^HIGH^ and 11.9% of genes were found differentially expressed between these two cell groups (Table S2). **B**. Pathway enrichment analysis based on differentially expressed genes between NNMT^LOW^NAMPT^LOW^ and NNMT^HIGH^NAMPT^HIGH^ cells. Venn diagram representation and list of terms commonly enriched in the upregulated genes of NNMT^LOW^NAMPT^LOW^ cell group found in each of the four datasets. Remaining terms and further analyses are listed in Table S3. **C**. CD44 downregulation is a marker of NAM-induced senescence of GB-PDC. Example of CD44 staining (R633) using FACS (left panel), and the corresponding quantification of CD44 IF signal per cell (right panel, unpaired *t* test with Welch’s correction, mean±SD, n = 3 independent biological samples). **D**. Multivariate analysis for the CD44, NAMPT and NNMT gene expression in TCGA-GB. CD44, NAMPT and NNMT are risk factors for progression free survival. Hazard Ratio = 2.51 (1.58-4). Statistical significance: * P ≤ 0.05.

Next, to identify specific cell surface markers of NAM-induced senescence in GB, we explored the commonly deregulated genes in NAM-treated GB-PDC and NNMT^LOW^NAMPT^LOW^ cells (Table S2). Of the 54 commonly deregulated genes, the hyaluronan receptor CD44 was consistently downregulated in both NAM-treated GB-PDC and the NNMT^LOW^NAMPT^LOW^ cells. In coherence with these findings, FACS analysis revealed a significant decrease in CD44 immunofluorescent signal per cell in NAM-treated GB-PDC compared to controls (Fig. 3C), identifying low CD44 expression as a potential marker of NAM-induced senescence in GB cells.

Finally, multivariate analysis combining NNMT, NAMPT and CD44 highlighted their expressions as a risk factors associated with worse progression-free survival of GB patient (Fig. 3D).

### Senescence induced by NAM is reversible

Since senescence is characterized by irreversible cell cycle arrest, NAM pretreatment is expected to prevent tumor formation by GB-PDC when engrafted into the brains of immunodeficient mice. To evaluate tumor development in response to NAM pretreatment, we used orthotopic xenografts of control and NAM-pretreated GB-PDC stably expressing luciferase (5706** and 6240**) for bioluminescent tumor monitoring (Fig. 4A). Although mice grafted with NAM-pretreated GB-PDC showed improved survival compared to controls, they eventually developed tumors (Fig. 4B). This unexpected result suggested either a subpopulation of NAM-treated GB-PDC was resistant to senescence or NAM-induced senescence was reversible. To distinguish between these possibilities, we monitored GB-PDC proliferation in response to NAM withdrawal compared to continuous NAM treatment. GB-PDC showed no increase in proliferation rates during continuous NAM treatment over several weeks. In contrast, after NAM withdrawal, the cells returned to control proliferation rates within 1–2 weeks (Fig. 4C, Fig. S4A). Clonality assays, in which single GB-PDC were seeded per well, confirmed that no cell resisted NAM-induced senescence. Senescent GB-PDC displayed the same clonal properties as control cells following NAM withdrawal (Fig. 4D). Next, we evaluated CD44 expression -identified as a marker of NAM-induced senescence- at the single-cell level in NAM-pretreated GB-PDC. FACS analysis revealed a complete restoration of CD44 expression after NAM withdrawal (Fig. 4E). We also assessed SA-βgal positivity and cell morphology, finding no differences between NAM-pretreated and control GB-PDC (Fig. S4B and C). Transcriptomic analysis showed that NAM-pretreated GB-PDC largely returned to their pre-senescent profiles. Only 1-4% of the ≈20.000 detected genes were differentially expressed between NAM-pretreated and control GB-PDC (Fig. 4F), with no commonly deregulated genes among the three NAM-pretreated GB-PDC (Table S4).

**Figure 4.**
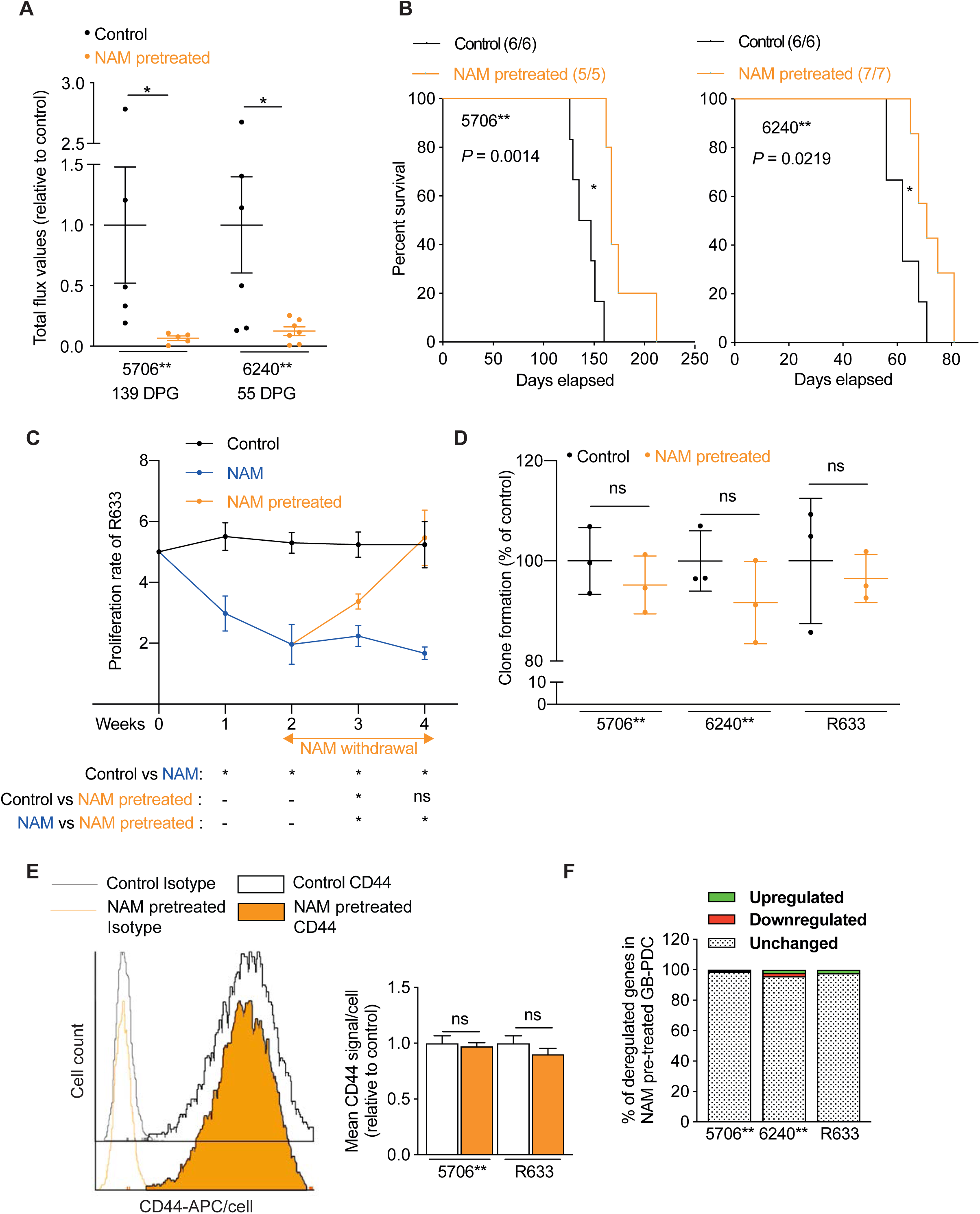
NAM-induced senescence is a reversible state. **A-B**. Mice grafted with NAM-pretreated GB-PDC have better survival. **A**. Bioluminescent analyses of tumor growth initiation by grafting either untreated (control) or NAM-pretreated GB-PDC transduced with a luciferase construct..Quantification of the bioluminescent signals. n = 5 (5706**) and n = 6–7 (6240**) mice per group. Mean ± SD. Unpaired *t* test with Welch’s correction. DPG: days post-graft. **B** Kaplan–Meier survival curves. 5706**: control n = 6 and NAM-pretreated n = 5; 6240**: control n = 6 and NAM-pretreated n = 7. Log-rank Mantel–Cox test. **C**. NAM withdrawal results in restoration of GB-PDC proliferation rate. Proliferation rate of control (week 1-4), NAM-treated (week 1-4) and NAM-pretreated (week 3-4) GB-PDC (R633). Unpaired *t* test with Welch’s correction, mean ± SD, n = 3 independent biological samples. **D**. NAM-pretreated cells have similar percentage of clone formation to control GB-PDC. Unpaired *t* test with Welch’s correction, mean ± SD, n = 3 independent biological samples. **E**. CD44 expression is restored in NAM-pretreated GB-PDC. Example of CD44 staining (R633) using FACS (left panel), and the corresponding quantification of mean CD44 IF signal per cell (right panel, unpaired *t* test with Welch’s correction, mean ± SD, n = 3 independent biological samples). **F**. NAM withdrawal restores GB-PDC transcriptomic profile. Gene fractions with upregulated, downregulated or unchanged expression following NAM withdrawal compared to control are highlighted in green, red or with a dot motif, respectively. Statistical significance: ns non-significant, * P ≤ 0.05.

These findings demonstrate that NAM-induced senescence is nearly fully reversible upon NAM withdrawal, restoring GB-PDC to pre-senescent state.

## Discussion

Emerging data highlight that metabolites not only participate in metabolic processes but also function as signaling molecules, profoundly influencing cell behavior (Baker and Rutter, 2023). In this context, we explored the non-canonical role of NAM and discovered that elevated intracellular NAM levels induce a cytostatic effect of GB cells corresponding to their entrance into an original state of senescence, exhibiting classical features of senescence while being totally reversible. Senescence, a complex cell state occurring in both physiological and pathological contexts,is characterized by significant alteration of cell properties. NAM-treated GB cells displayed hallmark features of this state, including reduced proliferation, changes in cell morphology, global epigenetic modifications, enhanced DNA damage, increased lysosomal activity, and reduced Lamin B1 expression.

Our study revealed that NAM-induced senescence in GB cells triggered an unexpected reprogramming of NAM metabolism, characterized by the downregulation of both NNMT and NAMPT, key regulators of DNA/histone methylation and NAD^+^ biosynthesis. Modeling this reprogramming at the single cell level in patients confirmed the link between low levels of NNMT and NAMPT and senescence. Our results regarding NAMPT are coherent with the documented decrease of its expression during aging (Khaidizar et al., 2021). In the same line, NNMT and NAMPT overexpression has been shown to correlate with a poor prognosis of GB patients (Jung et al., 2017; Lucena-Cacace et al., 2019; Palanichamy et al., 2017). Coherently, our findings showed improved survival in mice grafted with NAM-induced senescent GB cells.

NAM metabolism reprogramming is expected to influence several epigenetic marks. In our case, NAM-induced NNMT downregulation was accompanied by the upregulation of EZH1, resulting in higher global levels of the heterochromatin marker, H3K27me3. The state of H3K27me3 varies with different senescence contexts, being redistributed in replicative and oncogene-induced senescence and increased in stress-induced premature senescence (Paluvai et al., 2020). NAMPT downregulation combined with NAM intracellular accumulation in senescent GB-PDC is expected to affect the activity of NAD^+^-dependent enzymes activity, such as sirtuins. Sirtuins catalyze the deacetylation of many proteins, including histones. Among the different histone acetylation marks we explored, we found a consistent increase in H3K14ac levels in senescent GB-PDC. H3K14ac, implicated in genome stability and a marker of transcriptionally active chromatin, has been shown to regulate the expression of aging-related synaptic plasticity genes in mice (K. Wang et al., 2022). Overall, the epigenetic reprogramming observed in NAM-induced senescence in GB cells underscores the context-specific nature of these changes.

Mechanistically, we found that NAM-induced inhibition of NNMT was mediated by altered SOX2 protein expression. We showed that SOX2 protein binds to the promoter of NNMT and is downregulated following NAM treatment. Previous reports have observed senescence induction of GB cells following SOX2 downregulation (De Lope et al., 2019; Garros-Regulez et al., 2016a; Vinchure et al., 2021; Wu et al., 2020). SOX2 nuclear localization depends on several post-translational modifications, such as acetylation or methylation (Schaefer and Lengerke, 2020). Elevated acetylation of SOX2 due to inhibition or reduction of Sirtuin 1 resulted in nuclear exclusion and proteasomal degradation of the former (Baltus et al., 2009; Yoon et al., 2014). These data suggest that persistent NAM intracellular levels in NAM-treated GB-PDC could impair sirtuin activity, thus increasing acetylation levels of SOX2, leading to its degradation.

Despite being a well-characterized cell state, senescence lacks a single universal marker (Hernandez-Segura et al., 2018). Here, we identified downregulation of CD44 as a marker of NAM-induced senescence in GB. CD44 is a cell surface glycoprotein and receptor for hyaluronic acid, involved in cell-cell interactions, cell adhesion, and migration (Senbanjo and Chellaiah, 2017). In GB, CD44 is predominantly expressed in cancer stem-like cells and plays a pivotal role in GB cell invasiveness and resistance to chemotherapy (Inoue et al., 2023). In line with our findings, CD44 knockout has been recently reported to induce senescence of the U251MG GB cell line (Kolliopoulos et al., 2022). Moreover, NNMT has been reported to promote CD44 expression by decreasing H3K27 methylation in hepatocellular carcinoma (Li et al., 2019).

As expected, brain engraftment of control and NAM-pretreated PDC showed higher tumorigenic potential of control GB-PDC compared to their senescent counterpart, leading to a shorter survival of the respective grafted mice. Consistently, multivariate analysis combining NNMT, NAMPT, and CD44 highlighted their expression as risk factors predicting poorer progression-free survival in GB patients. Despite their improved survival, mice grafted with senescent GB-PDC eventually developed fatal tumors. This unexpected result contradicts the general assumption that senescence is characterized by an irreversible cell cycle arrest. Further experiments conducted at the single cell and bulk level showed that NAM-induced senescence is a reversible state. The irreversibility of senescence is a matter of debate in the literature. Accumulating data highlight the dynamic nature of senescence (Reimann et al., 2024). In some cases, the senescence program is executed to an extent that does not allow the stability of the terminal senescent state, despite cells expressing several senescence markers (Reimann et al., 2024). In contrast, other senescence programs result in a deep senescent state, from which cells can hardly recover despite withdrawal of the inducer of senescence (Reimann et al., 2024). In our case, NAM withdrawal resulted in restoration of GB-PDC properties to pre-senescent levels.

The role of senescence in cancer development is multifaceted. Senescence induction in pre-neoplastic cells, leading to a stable proliferation arrest, acts as a potent anti-tumor mechanism (Schmitt et al., 2022). Because senescence is extensively described as an irreversible cell growth arrest, its induction by anti-cancer therapies is expected to improve the clinical outcome of cancer patients. However, it has been shown that therapy-induced senescence could promote recurrence. Through their senescence-associated secretory phenotype (SASP), senescent cells boost the growth of their non-senescent counterparts (Schmitt et al., 2022). Besides the harmful effects of the SASP, therapy-induced senescence could constitute a strategy whereby tumor cells escape from the cytotoxic impact of therapy. Therapy-induced GB cell senescence is observed with the current temozolomide and irradiation therapies for GB patients (Aasland et al., 2019; Beltzig et al., 2022; Fletcher-Sananikone et al., 2021). Experimental data from non-brain tumors, such as breast and lung cancers, treated with cytotoxic drugs showed that senescence could be adopted in a transitory manner by cancer cells (Riviere-Cazaux et al., 2023). Upon exit from senescence, cancer cells re-entered into the cell cycle, showing that senescence participated in their escape from therapy (Riviere-Cazaux et al., 2023). Beyond therapy-induced senescence, senescent cancer cells have been identified in many untreated tumors, including GB (Ouchi et al., 2016; Salam et al., 2023), suggesting that the natural tumor progression relies on the presence of senescent cells (Lushchak et al 2023). Here, we discovered a novel facet of senescence in GB: metabolite-driven, completely reversible, and identifiable in patients’ tumors. We found that GB cells enter senescence in response to variations in their environment and emerge from it once their environment is restored. The role of this reversible senescence in the natural growth of the tumor remains to be explored.

In conclusion, we have discovered a novel non-canonical role of NAM: driving reversible senescence in GB cells. Our findings not only expand our understanding of the complex nature of senescence but also highlight the diversity of molecular changes associated with NAM-induced senescence. These findings suggest that the senescent state in GB cells can be a dynamic process, potentially influencing cancer cell plasticity and thereby therapeutic resistance.

## Acknowledgments

This work was supported by grants from Région Ile-de-France (MSS fellowship), Les Entreprises Contre Le Cancer GEFLUC Paris (Ile-de-France), La Fondation pour la Recherche Médicale – Equipe labelisée FRM and INCa PLBIO. We are grateful to Drs. D. Darmoul and F. Soualmia for scientific discussions.

## Material and Methods

### Biological material and NAM treatment

5706**, 6240** and R633 GB-PDC, derived from neurosurgical biopsy samples of distinct GB, were cultured in defined medium containing EGF and bFGF, as previously described (El-Habr et al., 2017; Rosenberg et al., 2017). 5706** and 6240** stably express luciferase.

Cell metabolic activity in presence of the different NAM (Sigma) concentrations was assessed using reduction in WST-1 (4-[3-(4-Iodophenyl)-2-(4-nitrophenyl)-2H-5-tetrazolio]- 1,3-benzene Disulfonate) to water-soluble formazan (Roche, France). Cells were seeded in 96-well plates at 2 × 10^4^ cells/well and treated with the different NAM concentrations at 37°C, 5% CO_2_. At the end of the incubation period, 10% (v/v) WST-1 was added to the culture media, and the cells were further cultured for 3 h. The absorbance was measured at 430 nm in a microplate reader (Expert Plus V1.4 ASYS).

Senescence was induced by treating GB-PDC with 30 mM of NAM for one (5706**) or two (6240** and R633) weeks. Two weeks after NAM withdrawal, GB-PDC were considered post-senescent. Control GB-PDC were treated with culture media.

### Cell proliferation, viability and morphology evaluation, and Senescence-associated β-galactosidase Staining

Proliferation: Trypan blue exclusion test was used to determine the number of viable cells (Trypan blue solution, Thermo-Fisher, 0.4% v/v, 3 min incubation at room temperature). Blue and white cells (dead and alive, respectively) were counted with the Countess automated cell counter (Thermo Fisher, France). For cell proliferation evaluation, cells were seeded in defined medium treated or not with NAM, and viable cell numbers were counted 1 week later. Viability: cell death was evaluated using the propidium iodide (PI) exclusion test.

Cells were incubated with PI (10 µg/10^6^ cells) for 10 min at 4 °C, and the percentage of cells containing PI was measured using FACS (ARIA II, BD Biosciences, France).

Morphology: cell morphology was assessed using FACS forward (for cell size) and side (for cell granularity) scatter (FSC and SSC respectively).

Clonality: control or NAM-pretreated GB-PDC were seeded at one cell/well (Nunc, 96-deep well plate, non-treated). The percentage of wells containing spheres was scored 2 weeks after cell seeding. At least 500 cells were analyzed for each culture.

For senescence-associated β-galactosidase staining, cells were smeared on SuperFrost slides (Fisher Scientific, France) and stained according to the manufacturer’s protocol (Cell Signaling Technology). The percentage of blue cells was determined using a Nikon inverted phase-contrast optical microscope.

### Metabolite measurement by mass spectrometry (MS)

Cells were harvested 96 h post-seeding (cell half-doubling time = 4.5, TG1, and 8 days, TG1-miR). Cell pellets were washed in PBS before freezing. For NAM and MNAM, cell samples (n = 6) were extracted and analyzed on the GC/MS and LC/MS/MS platforms of Metabolon, Inc. (Durham, NC, USA) as previously described (El-Habr et al., 2017; Reitman et al., 2011). For SAM and SAH in TG1 versus TG1-miR (n = 3), cell samples were extracted and analyzed with GC–MS/MS (300MS, Brüker) in the clinical chemistry laboratory at Necker Enfants Malades Hospital (Paris, France). For NAM in NAM-treated 5706** and 6240** (n = 1) cell samples were extracted with chloroform after acetone deproteinization and analyzed with Ultimate 3000 HPLC system (ThermoFisher Scientific) in the Metabolic and Cellular Biochemistry department at Bichat Hospital (Paris, France).

### Gene expression and ontology analyses

Total RNA was extracted using the Nucleospin RNA kit (Macherey-Nagel) according to the manufacturer’s instruction. cDNA was prepared using the QuantiTect Reverse Transcription Kit (Qiagen) according to manufacturer’s instructions. Expression profiles of TG1 and TG1-miR were determined using Affymetrix 1.0 Human Exon ST arrays in three independent cell passages as previously described (Bogeas et al., 2018).

Expression profiles of control, NAM-treated and NAM-pretreated 5706**, 6240** and R633 were determined in three independent cell passages using bulk RNA-sequencing analysis performed at Montpellier GenomiX plateform (MGX, Institut de Genomique Fonctionnelle, Montpellier, France). Strand-specific sequencing libraries were prepared from 500 ng total RNA using poly(A) capture of transcripts with the TruSeq Stranded mRNA Sample Preparation kit (Illumina). Libraries were quantified with the Fragment Analyzer (Standard Sensitivty NGS kit). Clusters were generated and sequencing was run on the Illumina NovaSeq 6000 using the NovaSeq Reagent Kits. Image analysis was performed using NovaSeq Control Software (Illumina), and base calling using the Real-Time Analysis 3 software (Illumina). Demultiplexing and fastq file generation were performed using bcl2fastq v2.20.0.422 software (Illumina). Quality control of sequencing data was obtained using FastQC v0.11.8 software. RNA-seq reads in fastq format were aligned and annotated to the Hg38 Homo sapiens reference genome (gtf file downloaded from University of California Santa Cruz (UCSC) website on 27/01/2020). The raw counts matrix was generated using featureCounts (v2.0.0) using default parameters, except for ‘reversely stranded’ option set at - s 2. Aligned data were quality-controlled using RSeQC (v2.6.4).

For differential gene expression, data was analyzed on R (v3.6.1). Raw counts matrix was first filtered to remove genes with low expression, keeping those with >10 reads in at least 3 samples, and then normalized using the “Relative Log Expression” (RLE) method (*calcNormFactors* function from edgeR v.3.26.8 R package (Robinson et al., 2010). Differential gene expression analysis was carried out on raw count data using exact test from edgeR R package. Significance levels was set at Benjamini-Hochberg (BH) adjusted p-value < 0.05. When comparing results from different GB-PDC, significance levels was set at p-value < 0.05.

RT-QPCR assays were performed using the LightCycler480 (Roche, France) and the SYBR Green PCR Core Reagents kit (Bimake.com). The thermal cycling conditions comprised an initial denaturation step at 94 °C for 5 min, and 40 cycles at 94 °C for 30 sec, 60 °C for 30 sec and 72 °C for 30 sec. Transcripts of the TBP gene encoding the TATA box-binding protein (a component of the DNA-binding protein complex TFIID) were quantified as an endogenous RNA control. Quantitative values were obtained from the cycle number (Cq value), according to the manufacturer’s manuals. Sequences of primers used for RT-QPCR are listed in Table S5.

### Single-cell RNA-sequencing analysis

#### Data was pre-processed and analyzed on R v3.6.1

Data acquisition and pre-processing. Four publicly available single-cell transcriptomes of GB cells were downloaded from the Broad Institute Single Cell Portal or GEO: Neftel_SMARTseq2, Neftel_10X, Yu and Wang_10X (Table S5). Malignant and normal cells were distinguished either according to the cell annotations provided by authors, or when absent, based on inference of copy-number variations (CNVs), a hallmark of malignant cells, as previously described (Saurty-Seerunghen et al., 2022) (Table S5, see Fig. S5 for Yu dataset and Sup Fig.1 of Saurty-Seerunghen et al., (Saurty-Seerunghen et al., 2022) for Neftel_10X dataset). Low-complexity transcriptomes were filtered out by authors for all datasets, except for the Neftel_10X dataset. For the latter, we considered cells with <150 genes, <250 UMI counts and >20% mitochondrial genes as being low-complexity transcriptomes or dead/dying cells and excluded them. We also excluded cells from Tumor105B-D from that dataset because they had lower numbers of genes and UMI counts detected than the other tumors. For all datasets, we retained only malignant cells from primary adult GB patients in further analyses. Therefore, we retained 4916 cells from twenty patients in the Neftel_SMARTseq2 dataset, 5797 cells from six patients in the Neftel_10X dataset, 2820 cells from eight patients in the Yu dataset, and 45361 cells from thirty-one patients in the Wang_10X dataset.

Cell grouping and differential gene expression analysis. GB cells, in which NNMT was detected, were first ranked based on NNMT expression levels and separated into two groups, NNMT^LOW^ and NNMT^HIGH^ based on the expression distribution’s mean (Fig. 3A, S3). This was then repeated for *NAMPT* gene. Next, identity of NNMT^LOW/HIGH^ and NAMPT^LOW/HIGH^ cells were compared to identify NNMT^LOW^NAMPT^LOW^ and NNMT^HIGH^NAMPT^HIGH^ GB cells for each dataset. Finally, genes differentially expressed between cell groups were identified following Mann-Whitney (Wilcoxon Rank Sum) test with p-values adjusted for multiple testing (Benjamini-Hochberg (BH), p-value < 0.01). Fold change (FC) for gene i was calculated as follows: FCi=xi−yi, where xi and yi are the log2 expression levels of gene i in conditions x and y, respectively. Only genes detected in at least 3% of GB cells were considered for this analysis. To avoid potential analytical bias due to scarcely detected genes, only genes detected in at least 3% of GB cells were considered for this analysis.

### Pathway and survival analysis

Pathway enrichment analyses on differentially-expressed genes were carried out on the enrichR website (https://maayanlab.cloud/Enrichr/). Pathways retained: Kyoto Encyclopedia of Genes and Genomes (KEGG).

A multivariate survival analysis of the combined expression of NNMT, NAMPT, and CD44 was conducted using the Tumor Online Prognostic Analysis Platform (ToPP) (http://biostatistics.online/topp/index.php). GB patients from TCGA (The Cancer Genome Atlas) were included in the analysis. The combined risk score for the three genes was calculated as follows: (0.136 * CD44) + (0.0166 * NAMPT) + (0.0556 * NNMT). Patients were then divided into high- and low-risk groups based on the median risk score.

### Protein expression analyses

Immunoblotting: cells were harvested, PBS washed and cell lysis was performed in 50 mM Tris–HCl pH 7.4 buffer containing 1% Triton X-100, 150 mM NaCl, 0.5 mM EGTA, 0.5 mM EDTA and anti-protease cocktail (Complete Protease inhibitor Cocktail Tablets, Roche, France). Protein extracts (30 µg) were separated by SDS-PAGE and transferred to Hybond-C Extra nitrocellulose membranes (GE Healthcare, USA). Details regarding primary and secondary antibodies are listed in Table S5. Signal detection was performed with the ECL+ chemiluminescence detection system (PerkinElmer, France). Densitometric analysis was achieved using ImageJ software.

Immunocytochemistry: cells were harvested, PBS washed, smeared on SuperFrost slides (Fischer Scientific, France), and fixed in PFA 4% for 15 min at room temperature (RT). Following fixation, cells were washed with PBS, and incubated for 30 min at RT in PBS containing 0.3% Triton X-100 and 5% BSA (Sigma). Details regarding primary and secondary antibodies are listed in Table S5. Immunostaining was analyzed with a fluorescent microscope equipped with an ApoTome module (Axioplan 2, Zeiss). Images were acquired on a digital camera using AxioVision 4.6 Software (Laboratory Imaging, Ltd) and prepared using Adobe Photoshop software (Adobe Systems, San Jose, CA). Immunofluorescent signals were analyzed with ZEN software (Zeiss).

CD44 expression analysis: single cell suspensions were prepared in PBS and stained with CD44 fluorophore-coupled antibody (Table S5). Cells were incubated at 4° C for 30 min under rotation. CD44 immunostaining was analyzed using FACS.

### ChIP-seq sample preparation and analysis

ChIP assays were performed using ChIP-IT Express Magnetic Chromatin Immunoprecipitation kit following the manufacturer’s protocol (Active motif, France) and 4 × 10^6^ cells per sample and per epitope. Briefly, cells were cross-linked in 0.5% formaldehyde/PBS for 10 min at room temperature and then treated with 0.125 M glycine in PBS pH 7.4 for 5 min at RT. Samples were subsequently washed twice with ice-cold PBS and once with ice-cold PBS supplemented with protease inhibitors cocktail prior to be lysed. Chromatin fragments ranging from 100 bp to 1000 bp were obtained by sonication. Chromatin was then incubated overnight at 4 °C on a rotor with anti-SOX2 (Table S5). The chromatin-antibody complexes were then washed, eluted and reverse cross-linked at 65 °C for 5 h. The eluted DNA was treated sequentially with Proteinase K and RNase A, and purified with the MinElute Reaction Cleanup Kit (Qiagen, #28204, France). The amount of DNA obtained was measured with a Qubit fluorometer (ThermoFisher, France). QPCR analysis was performed on total (input) and immunoprecipitated chromatin, and results normalized over the corresponding input signal. Enhanced representation of the regions of interest was compared to *TBP* promoter negative control. Sequences of all primers used for ChIP-QPCR are listed in Table S5.

### Intracranial xenografts

Animal experimentation was approved by Comité d’éthique en expérimentation animale Charles Darwin No. 5 (Protocol #5379). 5706** and 6240** stably expressing luciferase were used. 1.5 × 10^5^ of control or NAM-pretreated GB-PDC were injected stereotaxically into the striatum of anesthetized 8-10 weeks-old Nude female mice (Janvier Laboratories, France), and tumor formation and development monitored by bioluminescence imaging (Biospace Lab, M3 Vision software, France) as described previously (Saurty-Seerunghen et al., 2019).

### Statistical analyses

R or Prism 8.0 (GraphPad) softwares were used to generate plots and for statistical analyses. Significance level was set at p < 0.05. Each statistical test used is provided in the figure legends. When appropriate, the number of independent biological samples is indicated. Mean ± SD are shown.

---

**Figure S1, related to Figure 1.**
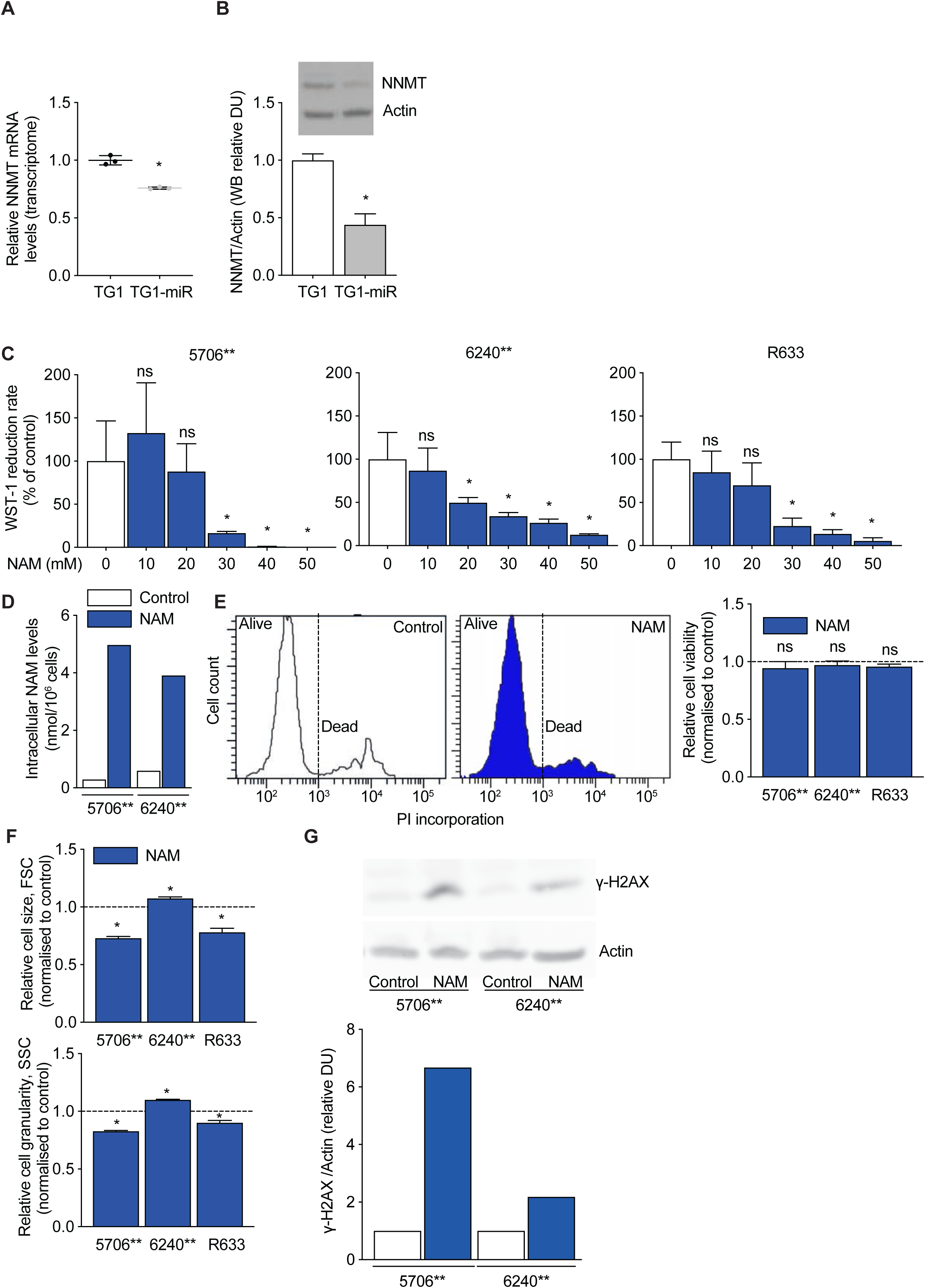
**A**. Decreased NNMT mRNA levels in TG1-miR compared to TG1. Microarray transcriptome analysis, unpaired *t* test with Welch’s correction, mean ± SD, n = 3 independent biological samples. **B**. Downregulation of NNMT protein levels in TG1-miR. Western blot analysis. NNMT: MW 30 kDa; Actin: MW 42 kDa; DU: densitometric unit. Unpaired *t* test with Welch’s correction, mean ± SD, n = 3 independent biological samples. **C**. NAM decreases the metabolic activity of GB-PDC. Cells were treated with NAM concentrations ranging from 10 to 50 mM, and reduction of WST-1 tetrazolium salt was measured as a readout of intracellular NAD(P)^+^/NAD(P)H ratios. Mann Whitney test, mean ± SD, n = 5 independent biological samples. **D**. NAM treatment leads to NAM intracellular accumulation in GB-PDC. HPLC analysis. **E**. NAM does not alter GB-PDC viability. Example of cell viability analysis (5706**) using FACS and propidium iodide (PI) incorporation (left panel), and the corresponding quantification of dead cell numbers (right panel, unpaired *t* test with Welch’s correction, mean±SD, n = 3 independent biological samples). **F**. NAM alters GB-PDC morphology. Quantification of cell size (Forward scatter, FSC, right panel) and granularity (Side scatter, SSC, left panel) using FACS. Unpaired *t* test with Welch’s correction, mean ± SD, n = 3 independent biological samples. **G**. Increase of *γ-*H2AX levels in NAM-treated GB-PDC compared to control. Western blot analysis. *γ-*H2AX: MW 17 kDa; Actin: MW 42 kDa; DU: densitometric unit. Statistical significance: ns non-significant, * P ≤ 0.05.

**Figure S2, related to Figure 2.**
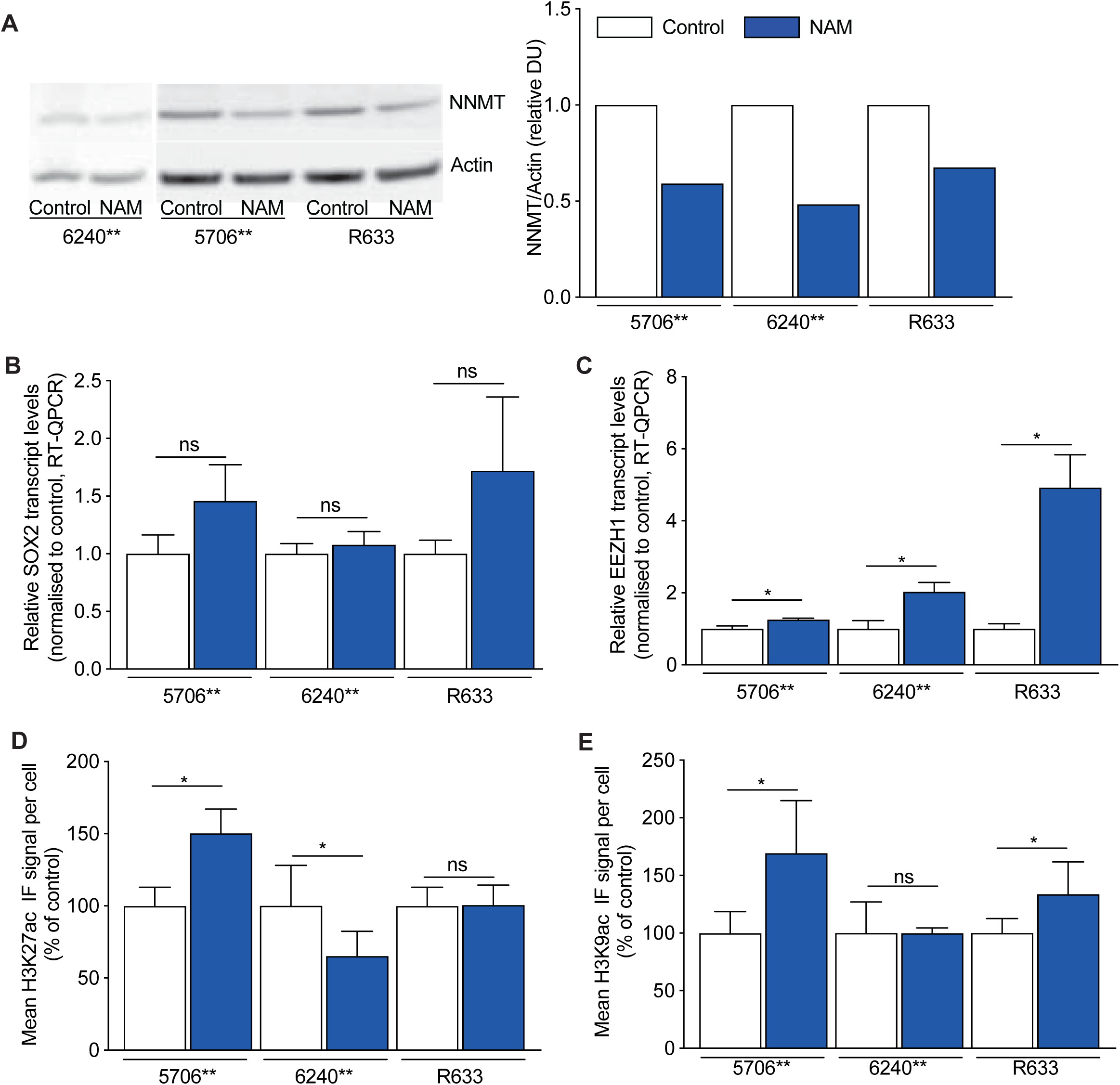
**A**. Downregulation of NNMT protein levels in NAM-treated GB-PDC compared to control. Western blot analysis. NNMT: MW 30 kDa; Actin: MW 42 kDa; DU: densitometric unit. **B**. NAM does not alter SOX2 transcript levels. RT-QPCR assays, unpaired *t* test with Welch’s correction, mean ± SD, n = 3 independent biological samples. **C**. Increased EZH1 mRNA levels in NAM-treated GB-PDC compared to control. RT-QPCR assays, unpaired *t* test with Welch’s correction, mean ± SD, n = 3 independent biological samples. **D-E**. H3K27ac (**D**) and H3K9ac (**E**) level variations in NAM-treated GB-PDC. Quantification of IF signal per cell for each histone mark. Unpaired *t* test with Welch’s correction, mean ± SD, n = 9 independent biological samples. Statistical significance: ns non-significant, * P ≤ 0.05.

**Figure S3, related to Figure 3.**
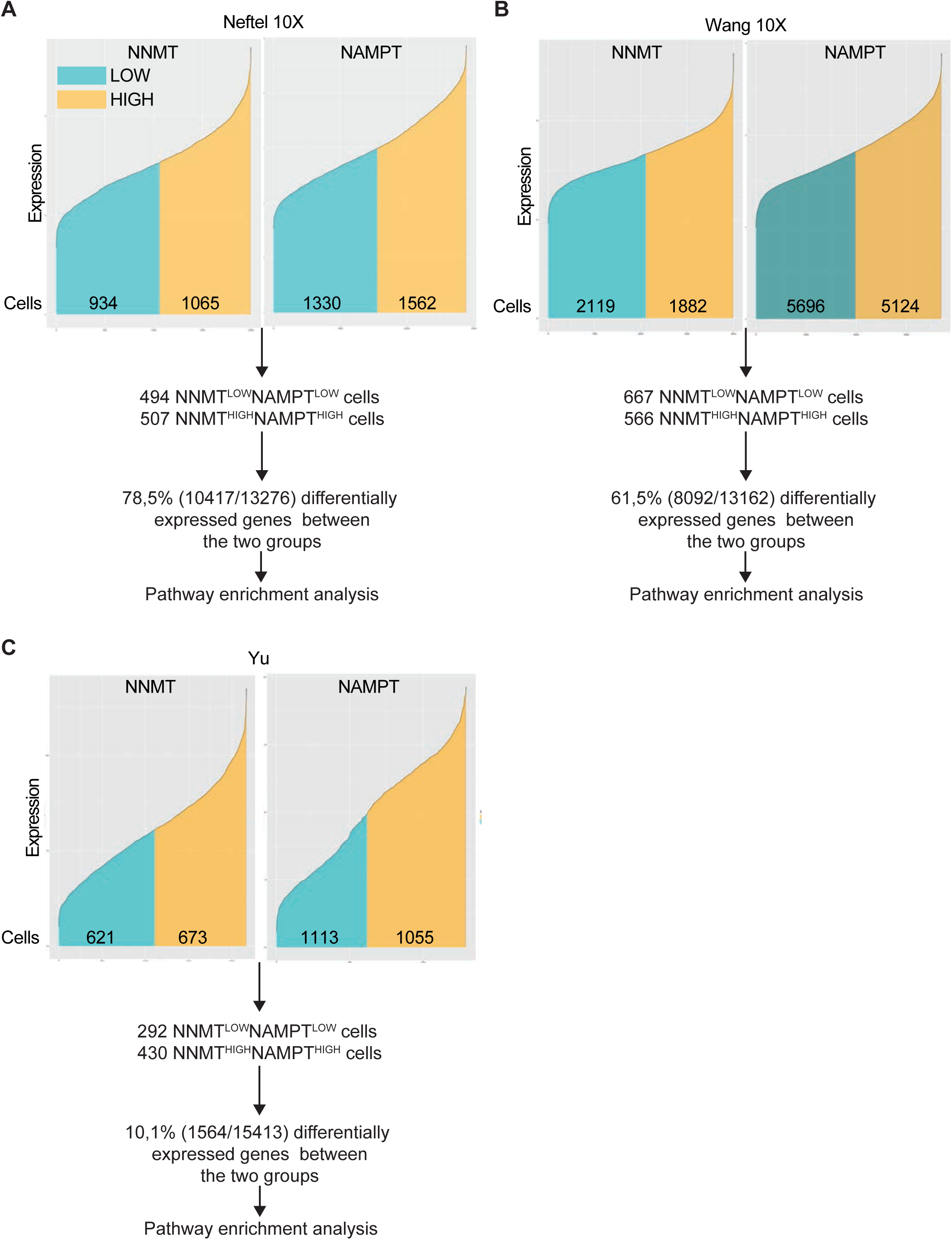
**A-C**. Scheme of the computational analysis of Neftel 10X (A), Wang 10X (B) and Yu datasets. Cells were split into two groups, NNMT^LOW^NAMPT^LOW^ and NNMT^HIGH^NAMPT^HIGH^, based on the mean expression of NNMT and NAMPT. The number of cells retained as NNMT^LOW^NAMPT^LOW^ and NNMT^HIGH^NAMPT^HIGH^ as well as the percentage of genes differentially expressed between these two cell groups are indicated. See text for further details.

**Figure S4, related to Figure 4.**
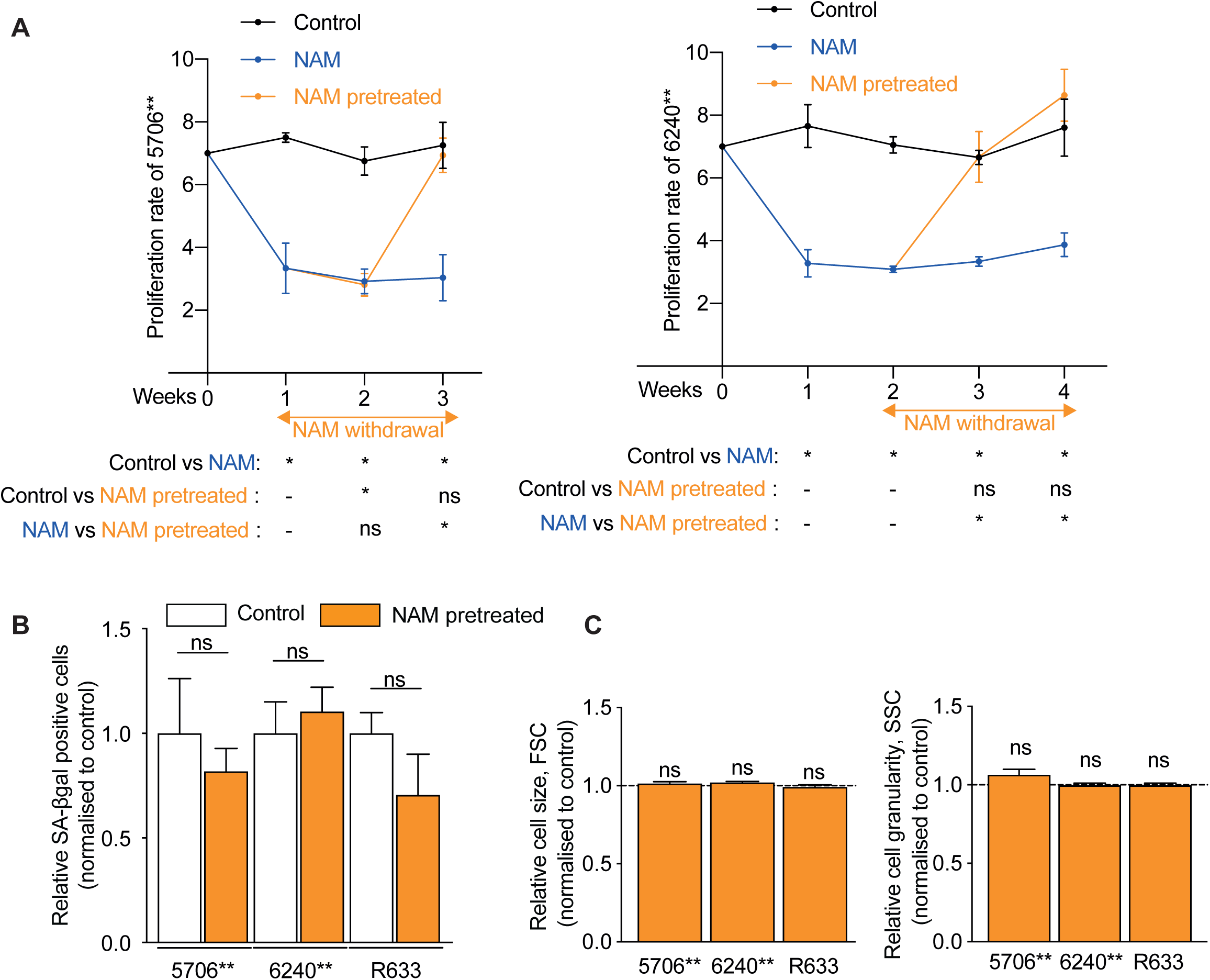
NAM withdrawal results in restoration of GB-PDC properties. **A**. Proliferation rate of control (week 1-4), NAM-treated (week 1-4) and NAM-pretreated (week 3-4) GB-PDC (5706**, left panel and 6240**, right panel). Unpaired *t* test with Welch’s correction, mean ± SD, n = 3 independent biological samples. **B**. Quantification of beta-galactosidase (SA-βgal) positive cells, unpaired *t* test with Welch’s correction, mean ± SD, n = 3 independent biological samples. **C**. Quantification of cell size (Forward scatter, FSC, left panel) and granularity (Side scatter, SSC, right panel) using FACS. Unpaired *t* test with Welch’s correction, mean ± SD, n = 3 independent biological samples. Statistical significance: ns non-significant, * P ≤ 0.05.

**Figure S5, related to Methods.** Identification of malignant and non-malignant cells in GB tumors from Yu dataset (Yu et al., 2020). **A**. Cell clustering based on GMM-based CNV predictions. Heatmap representation of CNV (copy number variations) predictions (**A1**). Potential malignancy status assigned, following cell clustering based on CNV predictions, highlighted on UMAP representation (**A2**). **B**. CNV predictions at canonical GB loci (Chr7 and 10). UMAP representation. **C**. Expression of marker genes of normal cell types. Pan-immune cells (PTPRC), macrophages (ITGAM, FCGR3A, CD14), microglia (CSF1R, TMEM119), T-cells (CD2, CD3D) and oligodendrocytes (MOG, MAG). UMAP representation. **D**. Cell clusters identified based on their repartition on UMAP plot. Hierarchical clustering followed by K-means clustering on UMAP components. **E**. Cell malignancy status assigned based on CNV status prediction, marker gene expression and clustering results. UMAP representation.

